# Diffusion MRI microstructure markers of changes in the human brain across the lifespan

**DOI:** 10.1101/2025.03.10.642447

**Authors:** Benjamin T. Newman, John D. Van Horn, T. Jason Druzgal

**Affiliations:** University of Virginia

## Abstract

Understanding how the brain develops, matures, ages, and declines is one of the fundamental questions facing neuroscience. Recent advances in diffusion MRI microstructure analysis have allowed for detailed descriptions of neuronal change in humans. However, it is essential that findings from these studies are appropriately contextualized to general age-related changes in the brain. This study uses 3-tissue constrained spherical deconvolution (3T-CSD) to examine the relationship between brain diffusion microstructure and chronological age. 3T-CSD is able to quantify signal fraction measurements at the voxel-wise level from three different tissue microenvironments found in the brain: extracellular free water, intracellular isotropic, and intracellular anisotropic. This study applies 3T-CSD analysis to the Nathanial Kline Institute’s Rockland cohort, a large-scale community sample of brain MRI data across the lifespan. Microstructural measurements were taken in a number of structures throughout the white matter, subcortical gray matter, and lobar cortical regions while additionally evaluating lateral differences in microstructural measurements. The general trajectory of signal fraction measurements was a positive relationship with age and extracellular signal fraction, a negative relationship between age and intracellular isotropic signal fraction, and an inverted U-shaped trajectory for the intracellular anisotropic signal fraction. In individual sub-areas these trends tended to still be present, with some notable exceptions. However there were large differences in 3T-CSD microstructure measurements between individual structures, including significant lateral differences between hemispheres for each of the subcortical gray matter structures and for each of the cortical regions. These results demonstrate that 3T-CSD is able to describe age-related change across the brain and lifespan. By using a healthy population cohort this study can be used as a point of comparison for 3T-CSD analysis of microstructure changes in the presence of pathology. Finally, the detailed analysis of lateralized ROI results can inform diffusion microstructure studies examining cortical and subcortical regions.

## Introduction

Throughout the human lifespan the structure of the brain changes dynamically in response to internal and external factors. A wealth of volumetric MRI studies have shown that early in life the brain dramatically increases in size while in older individuals the brain shows a remarkable decrease in volume(Driscoll et al., 2009; Fotenos et al., 2005; Good et al., 2001; Group, 2012; Pomponio et al., 2020; Raz et al., 2005). Two recent studies using extremely large numbers of subjects have firmly established a general trajectory for changes in brain volume, with peaks in volume occurring around 12 years of age (for subcortical GM, surface area, and cerebrum volume)(Bethlehem et al., 2022). However these trajectories show comparatively little movement following development (especially after puberty, but with the greatest rates of change occurring before 2 years of age)(Dima et al., 2022). This leads to a long and slightly downward plateau before decline begins in earnest in advanced age, with especially noticeable increases in ventricular volume and declines in cortical GM volume(Bethlehem et al., 2022; Dima et al., 2022). This lifespan trajectory is interwoven with complex processes during development and decline, such as hormone levels following the onset of puberty in early adolescence(Juraska & Willing, 2017) and changes in vascular health during aging(Jefferson et al., 2010). Concurrent with changes in gross anatomy are changes occurring at the cellular level, such as extensive synaptic pruning in development(Feinberg, 1982; Selemon, 2013) and excessive atrophy and cellular death in mild cognitive impairment and progressive decline(Stephan et al., 2012). Understanding the trajectory of these changes in the brain, as well as understanding what processes drive these changes, are important for research into neurological disease.

Recently, measures of cellular brain microstructure derived from diffusion MRI (dMRI) have proliferated in number and become more commonly applied in the analysis of development and pathology(Pasternak et al., 2009; Zhang et al., 2012). Each model has particularly defined functions applied to the raw diffusion signal, such as the separation of freely diffusing water from tissue in the bi-tensor free water elimination model(Pasternak et al., 2009) the ‘ball and stick’ model of isotropic and anisotropic diffusion (Assaf & Basser, 2005), or the detailed delineation of cellular somas from neurites (dendrites, axons, etc.) and extracellular water available with NODDI(Zhang et al., 2012). While these models have seen widespread application in recent years, most are still reliant on tensor-based diffusion models or require sufficiently detailed acquisitions to accurately quantify all of their output metrics. Our lab has recently developed a model of diffusion microstructure known as 3-tissue constrained spherical deconvolution (3T-CSD)(Newman et al., 2020). 3T-CSD relies on more complex models of diffusion signal calculated using spherical harmonics(Tournier et al., 2007) and is able to measure both isotropic and anisotropic intracellular microstructure and extracellular water compartments derived from single-shell clinical quality acquisitions with a high level of reliability and stability(Dhollander & Connelly, 2016; Newman et al., 2020). The ability of 3T-CSD and other quantitative diffusion models to measure multiple microstructure compartments provides a far greater level of detail at the voxel-wise level compared to well-established volumetric MRI and voxel-based morphometry measurements. These advantages mean that 3T-CSD is well-positioned to investigate changes in brain cellular microstructure across the lifespan. Within a specific age cohort, deviations from established age-related trajectories might indicate accelerated or slowed aging or development, depending on the context being studied. Microstructural measurements may have increased sensitivity by detecting changes in cellular architecture before they become apparent as changes in volume or density, particularly the signal from extracellular freely diffusing cerebrospinal fluid (CSF) infiltrating into brain tissue(Blair et al., 2019).

3T-CSD has several other advantages in examining changes in the brain across the lifespan. Importantly the intracellular isotropic component is flexible and nonspecific, and is intentionally not restricted to any specific cellular type or location within the brain. Changes in this signal fraction compartment are thought to arise from either neuronal soma or glial cell population changes. Voxels with the highest intracellular isotropic signal fraction occur in the typically defined gray matter areas (GM) in the cortex and subcortical structures. However nearly all voxels have a non-trivial contribution from each signal fraction compartment, including in axonal areas (so called ‘deep’ white matter (WM)) where neuronal somas are unlikely to be found(Khan et al., 2021; Mito et al., 2020). This makes the intra- and extracellular isotropic signal fraction compartments excellent markers for changes in axonal white matter areas. 3T-CSD can detect meaningful shifts from very high (∼90% or more) intracellular anisotropic signal fraction in healthy axonal areas toward increasing intra- and extracellular isotropic signal fraction measurements in early development(Pietsch et al., 2019) or in the presence of pathology(Khan et al., 2021). But this specific pathological context is not always available and as 3T-CSD begins to be applied to the study of broader conditions and developmental situations additional context is necessary to formulate hypotheses regarding the physiological change to the cellular microstructure that is observed in human studies. This context is especially necessary to interpret results from studies observing the relationship between cellular microstructure and variables with undetermined effects on neuronal physiology, such as pubertal hormones.

With the trajectory of volumetric MRI well established it is necessary to explore the cellular microstructure that underlies volumetric change with much greater detail, both with advanced dMRI models such as 3T-CSD and with detailed analysis of specific regions of the brain at multiple scales and between hemispheres. The goal of this study is to analyze a large number of subjects from a publicly available population cohort in order to provide a lifespan trajectory for 3T-CSD measurements of cellular microstructure from multiple areas of the brain. Establishing the relationship between chronological age and microstructural metrics will establish normal reference ranges and trajectories that are essential before investigating abnormal populations. This study will additionally investigate differences in cortical and subcortical areas and lateralized effects to inform future studies of significant regional and laterality differences that must be accounted for.

## Methods

### NKI/Rockland Cohort Data Acquisition

Data for this study was obtained from the Nathaniel Kline Institute’s Rockland Study (NKI/Rockland) cohort(Nooner et al., 2012). NKI/Rockland is a large-scale community sample of participants with ages across the lifespan gathered in Rockland County, New York. Rockland County was selected in part because its diverse ethnic and economic demographics resemble those of the United States as a whole, which aids generalizability to the broader population. A wide array of physiological, cognitive, genetic and neuroimaging assessments are collected and publicly released (available at: http://fcon_1000.projects.nitrc.org/indi/enhanced/index.html) (Nooner et al., 2012).

In this study the 409 subjects from the NKI/Rockland study that passed quality control and were publicly available at the time proceeded to analysis. These subjects ranged in age from 6-85 years-old (mean 42.67 ± 20.79 S.D.). There were 144 male and 265 female participants with average ages of 36.09 ± 21.22 S.D. and 46.25 ± 19.68 S.D., respectively. These ages were not equivalently distributed between sexes (Kolmogorov-Smirnov; D=0.276, p<0.001) which precluded performing analysis to compare trajectories between sexes.

### Diffusion image processing and analysis

dMRI data from 521 subjects was acquired using a Siemens MAGNETOM TrioTim 3T scanner with an isotropic voxel size of 2.0⨉2.0⨉2.0mm^3^, TE=85ms and TR=2400ms; 9 b=0s/mm^2^ images and 127 gradient directions at b=1500 s/mm^2^. These images were processed through an automated pipeline with several manual quality control steps as follows. Each diffusion image set was analyzed using SS3T-CSD(Dhollander & Connelly, 2016; Jeurissen et al., 2014) implemented in the open source software MRtrix and Mrtrix3Tissue(Dhollander & Connelly, 2016; Tournier et al., 2019). Several preprocessing steps utilized FSL(Jenkinson et al., 2012; Smith et al., 2004). Diffusion images were denoised (Veraart et al., 2016), corrected for Gibbs ringing(Kellner et al., 2016), susceptibility distortions(Smith et al., 2004), subject motion(Andersson et al., 2016), and eddy currents(Andersson & Sotiropoulos, 2016). All images were upsampled to of 1.3⨉1.3⨉1.3mm, and skull-stripping was performed using the Brain Extraction Tool (Jenkinson et al., 2012). Response functions were generated(Dhollander et al., 2016) from each tissue type and used to generate fiber orientation distributions (FODs)(Jeurissen et al., 2014). Signal fractions were calculated directly from the FODs using 3-tissue constrained spherical deconvolution (3T-CSD), a method that measures cellular microstructure within each voxel fitting into intracellular anisotropic (ICA, white matter-like), intracellular isotropic (ICI, gray matter-like), and extracellular isotropic (ECI, cerebrospinal fluid-like/Free Water) compartments(Newman et al., 2020). Whole brain measurements were calculated from native space signal fraction compartments. These metrics used a smaller whole brain mask that excluded the ventricles and subarachnoid space via a ECI threshold that restricted the voxels to only those containing a majority signal fraction from brain tissue compartments and not extracellular fluid.

White matter FODs were used to generate a cohort-specific FOD template image from 50 individuals between ages 32-38 years-old and each subject’s individual WM FOD image was registered to this template. Manual quality control of registration to the cohort-specific FOD template was performed by visual inspection, excluding any subjects with obvious distortions or extreme shears. Following this procedure 409 subjects remained in the study for analysis. An FOD template in stereotaxic space was created using the NTU-DSI-122 template (https://www.nitrc.org/projects/ntu-dsi-122/ ), extracting the equivalent single b-value shell and was registered to the cohort-specific template generated from this study(Hsu et al., 2015). This allowed for two separate atlases to be warped into the cohort template space to measure signal fraction averages within specific regions of interest.

### Regions of Interest

In order to cover a wide range of brain regions including cortex and deep white matter both the JHU-DTI based ICBM-DTI-81 white matter atlas (available as part of FSL, hereafter referred to as the JHU WM atlas) including 48 ROIs(Hua et al., 2008; Jenkinson et al., 2012; Mori et al., 2005; Wakana et al., 2007), and the Destrieux atlas including 164 ROIs(Destrieux et al., 2010) were warped into the study space. The cortical ROIs from the Destrieux atlas were summarized in a whole cortical ribbon ROI, as well as a cerebellum GM, and a total subcortical GM ROI. The 48 ROIs from the JHU WM atlas were also combined to provide a summary of the WM skeleton.

Subcortical GM ROIs were utilized from the Destrieux atlas, the nucleus accumbens, amygdala, caudate, hippocampus, putamen, and thalamus were included without alteration. The Destrieux atlas cortical ROIs were also separately combined into one of 8 lobar cortical regions per hemisphere as specified by the atlas and packaged in Freesurfer(Destrieux et al., 2010; Fischl, 2012). The cortical ROIs separately encompassed frontal, insular, limbic, motor, occipital, parietal, sensory, and temporal cortices.

### Statistical Approach

Signal fraction values from each of the three tissue compartments (ICA, ICI, and ECI) were averaged within each of the ROIs for use in analysis. The average values for the three tissue compartments were plotted against age for each of the 409 subjects. Total age-related change was examined both across these age groups and within each phase by the plotting of locally weighted scatterplot smoothing lines (loess) lines of best fit. This was performed in R using the default settings of a 2^nd^ order polynomial and span of 0.75 (meaning the localized slope reflects the closest 75% of data points). As a model-free way of displaying general trends loess lines were selected for descriptive purposes to show overall change in each signal fraction and facilitate between ROI comparisons. To provide more precise information on rates of change simple linear models were calculated predicting signal fraction from subject age and the slope of that relationship displayed for each ROI. For ease of comprehension as well as to delineate periods of particular interest across the lifespan (i.e. development, late age-related decline) we have divided the subject cohort into 4 different life phases: 1) developmental phase before 20 years of age, 2) early adulthood phase between 20-40 years of age, 3) late adulthood phase between 40-60 years of age, and 4) senescence phase greater than 60 years of age. This division allows for an overview of the entire lifespan as well as comparisons to studies which only feature cohorts within a particular age range (for example, development or advanced age).

The lateralized approach taken in this study for each of the subcortical and cortical ROIs allows for the comparison for each signal fraction between the left and right hemispheres. Measurements were taken from the left and right ROIs and compared across the lifespan using pairwise t-tests to compare each lateralized region within subjects. P-values were subsequently adjusted using a Benjamini & Hochberg correction for multiple comparisons.

## Results

### Whole brain shows developmental shift from ICI to ICA, followed by increasing ECI in aging

As each whole brain signal fraction was individually collected in an equivalent native space they are displayed, colored by age, as a ternary plot to illustrate change across the brain, across the lifespan (Fig. 1). It was observed that whole brain metrics begin with a higher percentage of signal from the ICI compartment early in development, then progressing toward a greater signal fraction contribution from ICA in adulthood. Signal fraction contribution from ECI does not appear to change greatly during this time. During aging however, ECI increases at the expense of both ICA and ICI as the brain declines and atrophy occurs.

**Figure 1:**
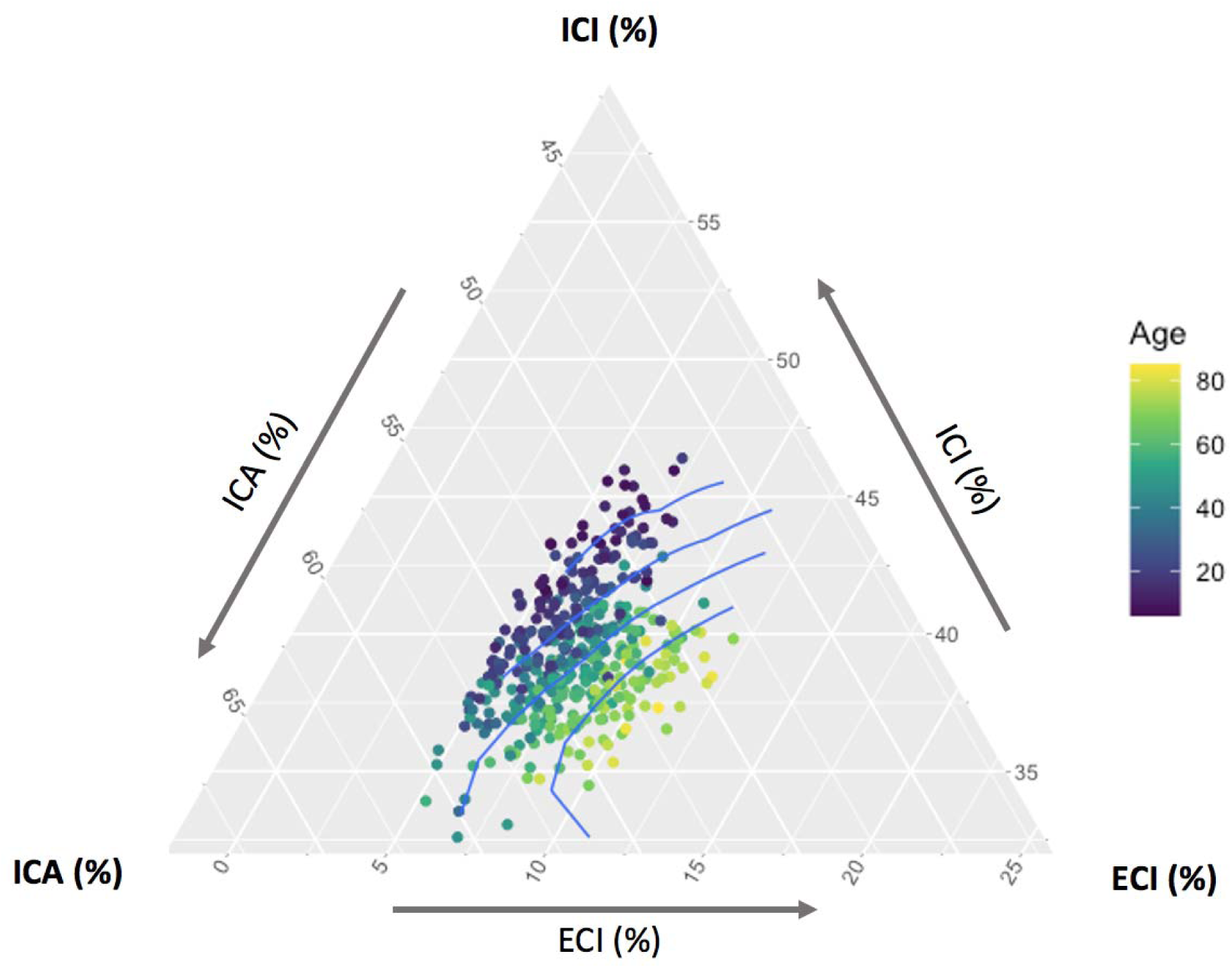
Ternary plot showing whole brain relationship between each of the 3T-CSD tissue compartments and subject age. Blue lines divide subjects into five 20%-tile groups based on age to illustrate change in signal fraction values across the lifespan. Whole brain tissue has the highest proportion of ICI signal fraction early in the lifespan, as development occurs the ICA signal fraction reaches its highest peak, and in later life the ECI signal fraction increases as the brain tissue declines.

### Large anatomical subregions show increasing ECI, decreasing ICI, and an inverted U-shaped ICA signal fraction across the lifespan

Loess lines of best fit (A) and the linear slope of change per year within each age group (B) are presented for the initial 4 generalized regions in Fig. 2. While the specific proportion of ICA, ICI, and ECI is different in each region, which is to be expected given that each region is largely defined by the predominance of a specific cell type, there is largely a consistent pattern of change in each across the lifespan. Matching the whole brain results, ECI showed low levels in development before 20 years of age, ECI even decreased in white matter areas covered by the JHU WM atlas and in subcortical GM structures. ECI remained largely flat in the early adulthood phase, between 20-40 years of age, but began to rise sharply during late adulthood between 40-60 years of age, with the sharpest increase occurring in the subcortical structures and cortical ribbon. In senescence greater than 60 years of age all major regions showed increasing levels of ECI reflecting widespread brain atrophy, loss of cellular integrity, and decline. The ICI signal fraction largely declined in all regions of the brain across the lifespan, though with important exceptions. The WM skeleton in the senescence phase showed a surprising *increase* in ICI signal fraction, especially considering that this is an area with very little ICI signal fraction. The cerebellum also remained relatively static in ICI signal fraction measurements until the senescence phase. The ICA signal fraction however, displays an inverted U shape across the lifespan, first rising greatly during development, slowing during the early adulthood phase, then increasing in decline during the early and senescence phases. The cerebellum is relatively consistent in ICA signal fraction in the development and late adulthood phases but otherwise each region is consistent in the trajectory of ICA measurements.

**Figure 2:**
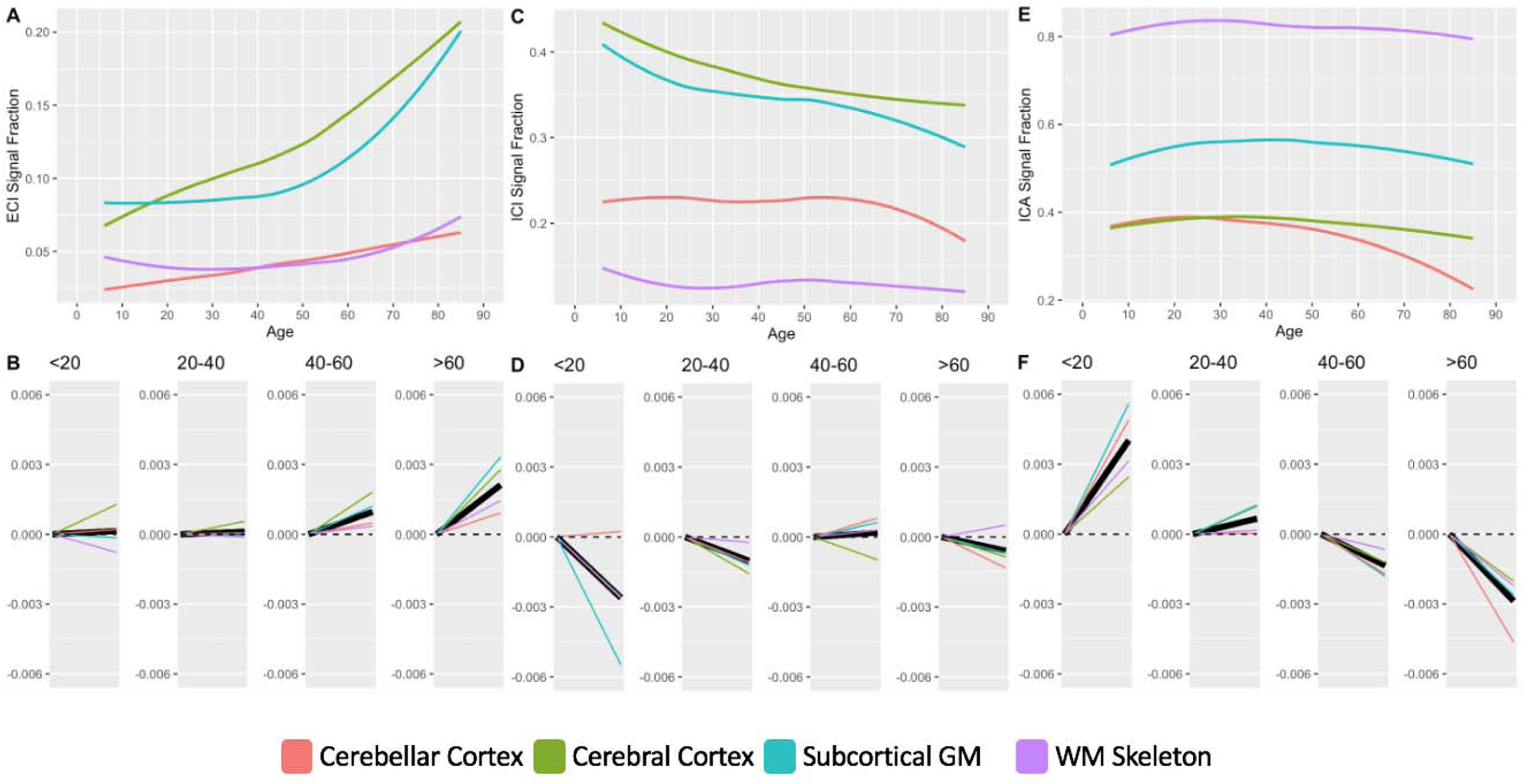
Charts displaying the lifespan trajectories of each 3T-CSD metric in 4 large anatomical brain subareas. The relationship between age and signal fraction is displayed either across the whole lifespan (A, C, & E for ECI, ICI, & ICA respectively) or as the slope of the linear relationship during a limited age range (B, D, & F for ECI, ICI, & ICA respectively). Overall, signal fraction compartment trajectories were relatively consistent between brain subareas, with a positive relationship between ECI and age, a negative relationship between ICI and age, and a positive relationship between ICA and age in the first two life phases, followed by a negative relationship between ICA and age in the later two life phases. There were however several interesting exceptions to this general trend. Within the WM skeleton there was a negative relationship between age and ECI signal fraction in the developmental phase, likely due to increased myelination reducing the extracellular space available for freely diffusing water. On the opposite end of the lifespan, in the senescence phase the WM skeleton displays an increase in ICI signal fraction.

### Subcortical GM ROIs typically follow whole brain trajectory but differ greatly from each other

Proceeding to a greater level of detail in different components of the subcortical GM, were independently measured in each hemisphere and plotted similarly to the larger anatomical areas (Fig. 3). A majority of the ROIs in this sample followed the general pattern observed for the area as a whole, but several important deviations can be observed, both in absolute differences in tissue composition as well as different trajectories across the lifespan. For example, the caudate nucleus early in life has a relatively low ECI signal fraction closer to the more internally located nucleus accumbens, but by middle age this structure contains a higher ECI signal fraction than the hippocampus, and the rate of increase then continues to accelerate until the structure has the highest average signal fraction of any subcortical structure late in life. Other structures demonstrate substantial deviations from the trajectory of the subcortex as a whole, most notably the putamen, which appears largely resistant to large age-related increases in ECI signal fraction, but in early development also the amygdala and hippocampus which decrease in ECI signal fraction more akin to the WM skeletal trajectory than the cortical trajectory. The basal ganglia appear to have the least ECI signal fraction and have matching trajectories, but the ventral striatum nucleus accumbens and the dorsal striatum putamen show different trajectories in the other signal fraction compartments indicating different responses to age-related change.

**Figure 3:**
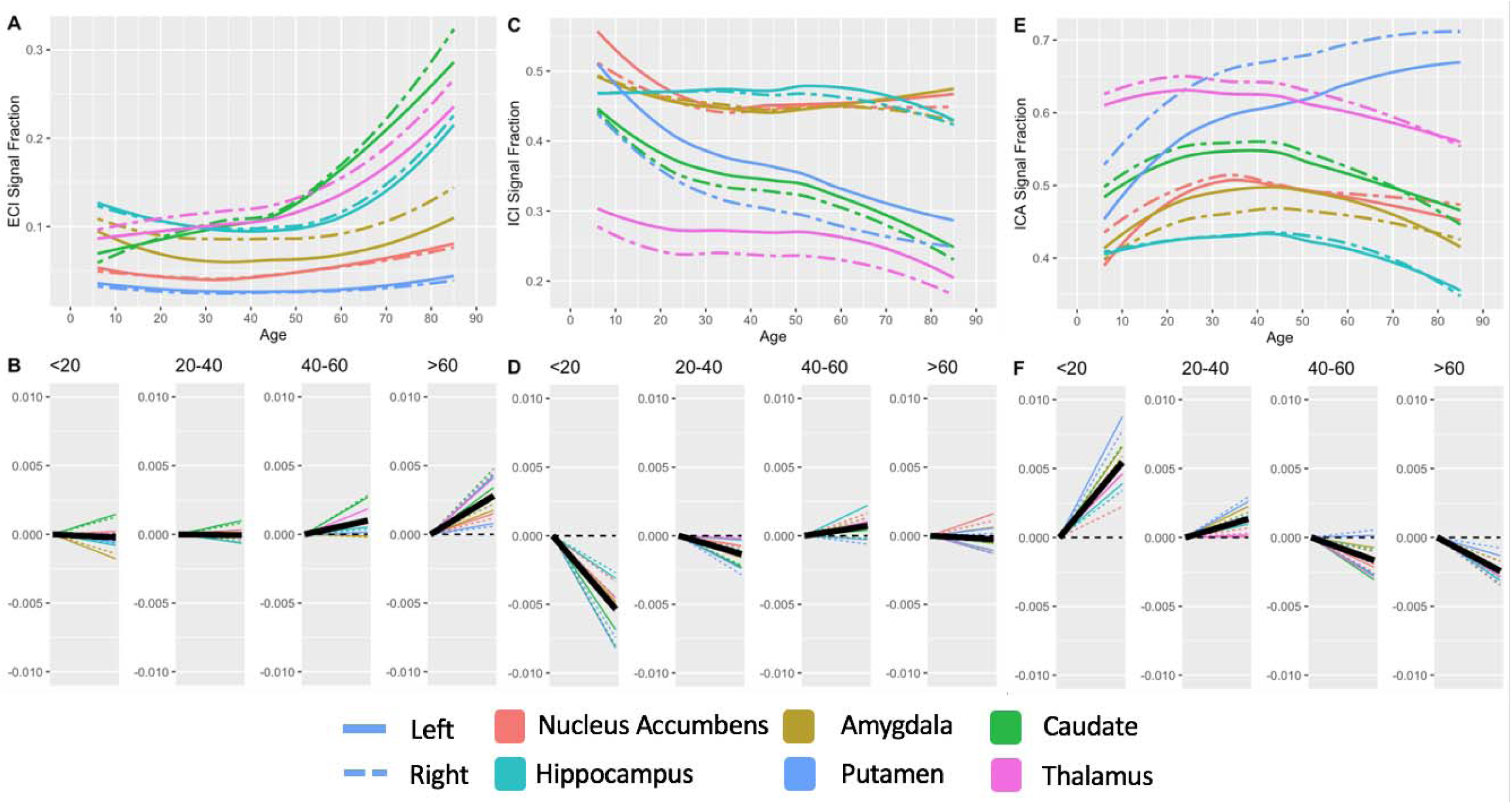
Charts displaying the lifespan trajectories of each 3T-CSD metric in 6 subcortical gray matter structures, including both left and right structures. The relationship between age and signal fraction is displayed either across the whole lifespan (A, C, & E for ECI, ICI, & ICA respectively) or as the slope of the linear relationship during a limited age range (B, D, & F for ECI, ICI, & ICA respectively). Trajectories largely follow the pattern established for the whole subcortical GM structure in Fig. 2, however some notable deviations: for the ECI signal fraction both amygdala, hippocampal, and amygdala drive the slight decrease in ECI signal fraction before 20 years of age, while caudate and thalamus surprisingly show steep increases during this time period and the hippocampus continues decreasing until reaching the late adulthood phase. In the ICI tissue compartment the thalamus, caudate, and putamen follow the whole subcortical trajectory, but the amygdala, nucleus accumbens, and to a near degree the hippocampus, are more resistant to age-related decline. In the ICA tissue compartment the putamen is likewise resilient to age-related decline.

While the trajectory of each subarea largely appears more or less extreme in the ECI signal fraction, in the ICI and ICA signal fraction compartments some of the subareas deviate entirely from the established trajectory. The amygdala, nucleus accumbens, and to a near degree the hippocampus, are more resistant to age-related decline and do not display substantial ICI signal fraction decrease in the early adulthood and late adulthood phases. These three regions approach the highest age included in this study with approximately 15% points higher (a greater than 50% increase over) ICI signal fraction than the putamen and caudate which begin development at a reasonably similar ICI signal fraction. The hippocampus follows a trajectory more akin to the cerebral cortex, but this is not relevant for the deep internal structure of the amygdala and nucleus accumbens. While each of the subcortical GM structures shows a wide range of ICA signal fraction compositions, the largest outlier is the putamen, which shows an increase in ICA signal fraction throughout the lifespan even as every other region begins to decline in the early and senescence phases. Toward the end of the lifespan the putamen even approaches within 10% points of the ICA signal fraction within the WM skeleton.

### Subcortical GM show significant and consistent lateralized differences

Results from lateralized comparisons are presented in Table 1. There was a high degree of lateralization in nearly all signal fractions with 15 out of 18 possible signal fraction/ROI combinations showing a significant difference between right and left hemispheres after correction for multiple comparisons. The ECI signal fraction was significantly higher in the right hemisphere in 4 of the 6 ROIs, as was the ICA signal fraction. The ICI signal fraction was higher in each left hemisphere ROI and was significantly so in all but the amygdala. The putamen, thalamus, and amygdala were observed to have the greatest laterality (at least one signal fraction compartment with a mean difference greater than 2%). The Hippocampus displayed the smallest average difference between left and right hemispheres, but was still significantly different for 2 of the 3 signal fractions, displaying how prevalent lateralized differences in structure are in the brain.

**Table 1:** Lateral differences between subcortical GM structures across the lifespan. Pairwise t-tests were used to compare each lateralized region within subjects and p-values were adjusted using a Benjamini & Hochberg correction for multiple comparisons. The ICI signal fraction was higher in each left hemisphere ROI and was significantly so in all but the amygdala. The putamen, thalamus, and amygdala were observed to have the greatest laterality (at least one signal fraction compartment with a mean difference greater than 2%).

| ROI | Tissue Type | Higher Signal Fraction | Mean Difference | t-value | Adjusted p-value |  |
| --- | --- | --- | --- | --- | --- | --- |
| **N. Accumbens** | ECI | *Left* | 0.0005 | 0.359 | 0.720 | n.s. |
|  | ICI | *Left* | 0.0106 | 2.634 | 0.012 | * |
|  | ICA | *Right* | 0.0100 | 2.139 | 0.040 | * |
| **Amygdala** | ECI | *Right* | 0.0232 | 19.596 | <0.001 | *** |
|  | ICI | *Left* | 0.0010 | 0.554 | 0.614 | n.s. |
|  | ICA | *Left* | 0.0226 | 11.264 | <0.001 | *** |
| **Caudate** | ECI | *Right* | 0.0049 | 3.073 | 0.003 | ** |
|  | ICI | *Left* | 0.0156 | 10.608 | <0.001 | *** |
|  | ICA | *Right* | 0.0106 | 7.158 | <0.001 | *** |
| **Hippocampus** | ECI | *Right* | 0.0029 | 2.517 | 0.016 | * |
|  | ICI | *Left* | 0.0066 | 3.995 | <0.001 | *** |
|  | ICA | *Right* | 0.0031 | 1.768 | 0.088 | n.s. |
| **Thalamus** | ECI | *Right* | 0.0169 | 21.483 | <0.001 | *** |
|  | ICI | *Left* | 0.0324 | 32.87 | <0.001 | *** |
|  | ICA | *Right* | 0.0155 | 12.574 | <0.001 | *** |
| **Putamen** | ECI | *Left* | 0.0021 | 5.632 | <0.001 | *** |
|  | ICI | *Left* | 0.0578 | 22.783 | <0.001 | *** |
|  | ICA | *Right* | 0.0598 | 22.542 | <0.001 | *** |

### Lobar cortical regions show similar trajectories across the lifespan but different absolute measurements and inconsistent but significant lateral differences

The cerebral cortex was then subdivided into 8 regions of cortex separated by left and right hemisphere as defined by the Destrieux atlas and measurements were made as before (Fig. 4). The signal fraction measurements in these regions generally followed the overall region trajectory displayed in Figure 3 but as with the subcortical GM there were some exceptions. The ECI signal fraction increased in every ROI in both the early and senescence phases. In the ICI and ICA signal fractions the insular and limbic cortices displayed markedly higher solid tissue values than other cortical regions while still following relatively similar trajectories. There was a significant degree of hemispheric laterality between the left and right sides of the brain (Table 2), with little consistent differences between the signal fraction compartments.

**Figure 4:**
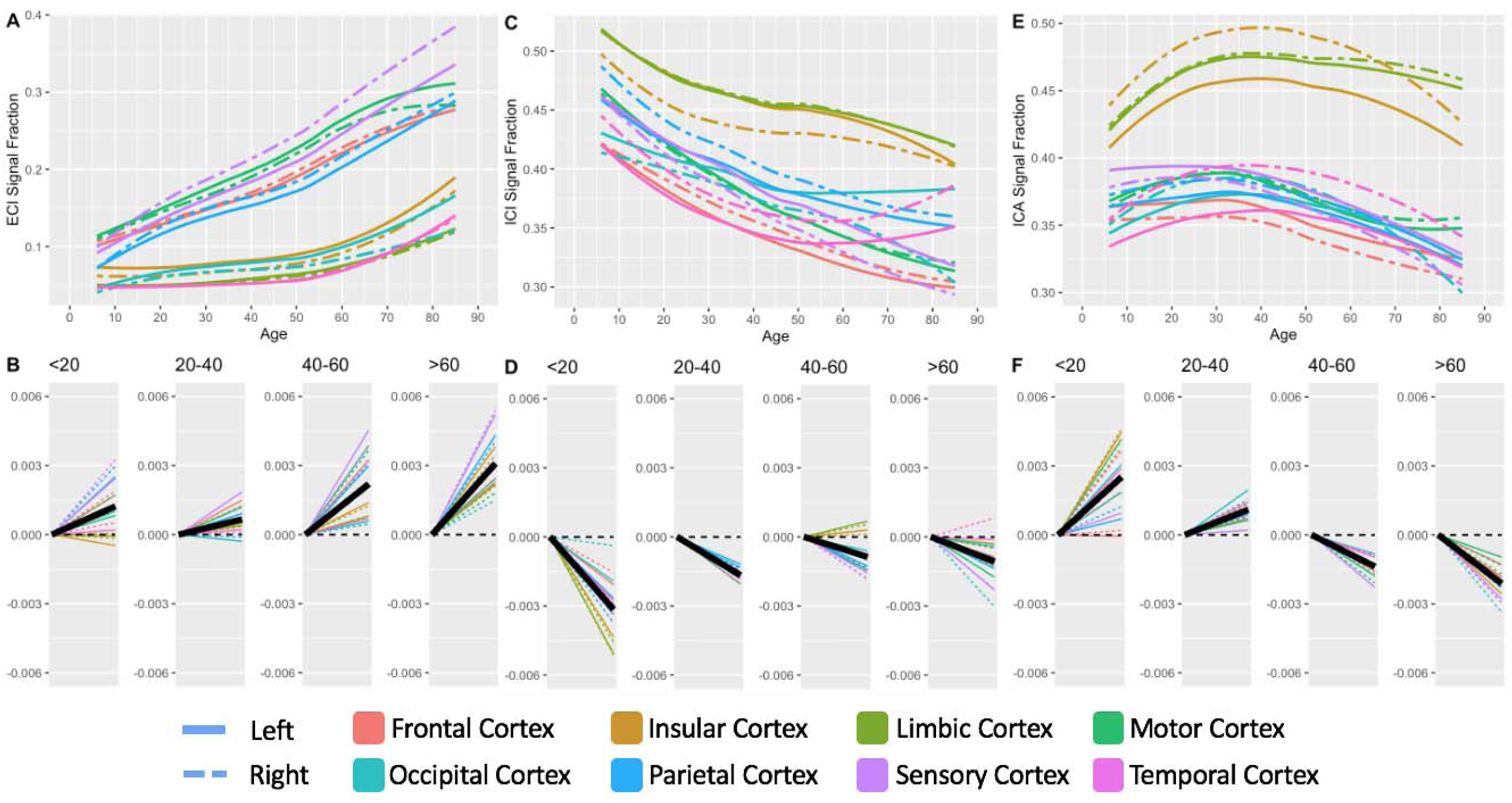
Charts displaying the lifespan trajectories of each 3T-CSD metric in 8 cortical ROIs. The relationship between age and signal fraction is displayed either across the whole lifespan (A, C, & E for ECI, ICI, & ICA respectively) or as the slope of the linear relationship during a limited age range (B, D, & F for ECI, ICI, & ICA respectively). Several deviations from the average trajectory occur in the middle age phases in the cortex in the insular and limbic cortices in the ICI signal fraction but otherwise declines across the lifespan. The frontal cortex in the developmental age phase shows little to slightly positive change in the ICA signal fraction but otherwise follows a consistent upward trajectory during the initial two life phases followed by decreases in the next two life phases.

**Table 2:** Lateral differences between cortical GM regions across the lifespan. Pairwise t-tests were used to compare each lateralized region within subjects and p-values were adjusted using a Benjamini & Hochberg correction for multiple comparisons. All cortical regions showed some significant degree of lateralization with the insular and temporal cortices each having a mean difference of greater than 2% ICA and ICI signal fraction measurements between left and right hemispheres. Conversely the limbic and motor cortices each had a mean difference of less than 0.5% excepting the ECI signal fraction in the motor cortex. Interestingly, as opposed to the subcortical areas which demonstrated relatively consistent right/left biases for the signal fraction compartments, there was a more balanced distribution of right/left biases for each of the signal fractions.

| ROI | Tissue Type | Higher Signal Fraction | Mean Difference | t-value | Adjusted p-value |  |
| --- | --- | --- | --- | --- | --- | --- |
| **Frontal Cortex** | ECI | *Right* | 0.0052 | 6.861 | <0.001 | *** |
|  | ICI | *Right* | 0.0088 | 9.783 | <0.001 | *** |
|  | ICA | *Left* | 0.0111 | 12.376 | <0.001 | *** |
| **Insular Cortex** | ECI | *Left* | 0.0121 | 16.632 | <0.001 | *** |
|  | ICI | *Left* | 0.0207 | 15.656 | <0.001 | *** |
|  | ICA | *Right* | 0.0339 | 30.966 | <0.001 | *** |
| **Limbic Cortex** | ECI | *Left* | 0.0021 | 4.365 | <0.001 | *** |
|  | ICI | *Right* | 0.0012 | 1.896 | 0.067 | n.s. |
|  | ICA | *Right* | 0.0036 | 4.665 | <0.001 | *** |
| **Motor Cortex** | ECI | *Left* | 0.0088 | 5.579 | <0.001 | *** |
|  | ICI | *Left* | 0.0009 | 0.742 | 0.4583 | n.s. |
|  | ICA | *Right* | 0.0021 | 1.425 | 0.1692 | n.s. |
| **Occipital Cortex** | ECI | *Left* | 0.0124 | 14.573 | <0.001 | *** |
|  | ICI | *Left* | 0.0196 | 7.591 | <0.001 | *** |
|  | ICA | *Right* | 0.0078 | 4.227 | <0.001 | *** |
| **Parietal Cortex** | ECI | *Right* | 0.0116 | 10.953 | <0.001 | *** |
|  | ICI | *Right* | 0.0137 | 15.809 | <0.001 | *** |
|  | ICA | *Right* | 0.0089 | 12.574 | <0.001 | *** |
| **Sensory Cortex** | ECI | *Right* | 0.0312 | 16.461 | <0.001 | *** |
|  | ICI | *Left* | 0.0186 | 13.752 | <0.001 | *** |
|  | ICA | *Left* | 0.0124 | 8.283 | <0.001 | *** |
| **Temporal Cortex** | ECI | *Right* | 0.0008 | 1.232 | 0.2282 | n.s. |
|  | ICI | *Right* | 0.0209 | 14.775 | <0.001 | *** |
|  | ICA | *Right* | 0.0308 | 19.764 | <0.001 | *** |

## Discussion

This study examined a large population cohort using an advanced diffusion microstructure analysis technique to describe the relationship between chronological age and changes in brain cellular structure. The slope of these changes was steeper during the developmental phase before 20 years of age and in the senescence phase after 60 years of age with more gradual changes occurring between 20 and 60 years of age in the middle of the adult phases of the lifespan. The pattern established at the whole brain level, a shift toward increasing ICA signal fraction in the first half of the lifespan largely coupled with decreasing ICI signal fraction (as they sum to 1, any increase/decrease in one signal fraction compartment must be coupled with equivalent increases/decreases in the others); subsequently gives way to a decline pattern later in life characterized by increasing ECI signal fraction coupled with decreasing ICA signal fraction.

When the WM skeleton, cerebral cortex, cerebellular cortex, and subcortical GM were parcellated from the cohort it was observed that the ECI signal fraction increased precipitously with increasing age after 40 in all 4 ROIs but during development was stable in the subcortical GM and cerebellum, declined in the WM skeleton, and increased in the cortex. Despite three of the ROIs being thought of as predominantly GM areas there was observed a widespread negative correlation with age throughout the lifespan. During the developmental phase the ICA signal fraction increased dramatically in each of the 4 ROIs, matching the well-established trajectories observed in volumetric data from this time period(Lebel & Beaulieu, 2011; Lebel & Deoni, 2018). Interestingly, the increased axonal innervation of the cortex during development can be observed in this trajectory as well as in the subcortical GM structures. This axonal innervation has been suggested to shift the WM/GM boundary line in volumetric studies(Natu et al., 2019) and this study certainly supports the cortex and subcortical GM structures becoming more similar to the WM skeleton in microstructural profile as development progresses. An interesting exception to this trend is in the frontal cortex, which has a flat slope for the relationship between ICA signal fraction and age during the developmental phase. This study also showed far more dramatic changes throughout the lifespan compared to large-scale volumetric MRI studies(Bethlehem et al., 2022; Dima et al., 2022). Change in brain metrics in this study was not solely confined to early development and late senescence, and there was very large and distinct trajectories for each tissue compartment across different ROIs, suggesting that age-related change is a highly complex process with different, distributed effects in different brain structures.

Laterality was examined and was found to have a highly significant effect across multiple regions but this was typically without a consistent direction. The putamen, thalamus, amygdala, insular cortex, and temporal cortex were observed to have the greatest laterality (at least one signal fraction compartment with a mean difference greater than 2%). The Hippocampus displayed the smallest average difference between left and right hemispheres, but the ECI and ICI signal fraction compartments were still significantly different, recapitulating previous work that found a significantly increased ECI signal fraction in the right hippocampus(Newman et al., 2020).

Other advanced dMRI models have examined the relationship between their quantitative measurements and age across the lifespan. In a recent study NODDI outputs of isotropic volume fraction, roughly comparable to 3T-CSD’s ECI signal fraction, increased throughout the lifespan across the whole brain. Meanwhile intracellular volume fraction displayed an inverted U shaped pattern similar to 3T-CSD’s companion ICA signal fraction(Beck et al., 2021). Another NODDI study however found conflicting results using orientation dispersion index, which does not have a direct 3T-CSD parallel, with results from Nazeri et al., finding orientation dispersion decreased across the lifespan in 4 cortical lobe ROIs, and Beck et al., finding that orientation dispersion increased across the whole brain(Beck et al., 2021; Nazeri et al., 2015). The current study found much more complicated relationships between many of the cortical lobes and age across the lifespan, and while this study did not directly compare results from different quantitative dMRI models, it is notable that NODDI and other advanced models did not perform better than traditional DTI metrics at predicting brain age(Beck et al., 2021).

There may be an expressed relationship in this study between signal fraction (particularly the relative proportion of ICA and ICI signal fractions) and neuronal cellular density as determined by histology. In particular the limbic cortex is known as having a particularly low neuronal density in humans(Semendeferi et al., 1998) and in this study was surprisingly found to have the highest level of ICI signal fraction (along with the insular cortex) throughout the lifespan. This suggests that a relatively low density of neurons may not cause a high ECI signal fraction but will instead be represented as an overly high ICI signal fraction, perhaps due to increased glial cell population or larger levels of intracellular space in comparison to areas of high neuronal and dendritic density, such as the cerebellum. This ROI did subsequently follow similar trajectories to other ROIs with differing levels of neuronal density and arrangement, but it regardless may hold promise as an effect replicable in pathological conditions affecting neuronal density. Decreased neuronal density in the prefrontal cortex has been implicated in major depressive and bipolar disorders, suggesting 3T-CSD may be a potential mechanism for tracking alterations in these patients(Cotter et al., 2001; Rajkowska et al., 2001).

Other disorders may result from increased neuronal density including schizophrenia which may present a different signal fraction profile that is able to be detected using 3T-CSD(Benes et al., 1986; Selemon et al., 1995). The developmental component of schizophrenia may allow for early detection if adolescents begin deviating from age-related trajectories similar to those established in this study. Another dMRI microstructural analysis technique, termed free water elimination, has suggested that increased extracellular water may also contribute to the development of schizophrenia(Pasternak et al., 2012), however this method does not measure both extracellular and isotropic intracellular tissue compartments and thus makes an imperfect point of comparison to the 3T-CSD technique used here.

A known feature of the aging brain that was not accounted for in this study is the development of white matter hyperintensities (WMH). Previous work has demonstrated that developed and developing WMH are detectable using 3T-CSD and have increased ECI and ICI signal fraction with correspondingly reduced ICA signal fraction compared to normal appearing areas of the WM skeleton(Mito et al., 2020). The presence of WMH not accounted for in this study for several reasons, including that the location of the WMH in the brain was largely determinative of its microstructural properties, with WMH located proximal to the ventricles having a greater amount of ECI signal fraction than WMH located within the deep WM(Mito et al., 2020). Additionally, there is evidence that 3T-CSD is sensitive to developing WMH that subsequently appear after cerebral infarction(Khan et al., 2021). This created a situation where removing tissue identified via independent means as a WMH might contribute noise to estimates of subjects in the senescence phase and would be difficult to account for in the whole lifespan trajectories. Another feature not accounted for was subject sex due to the imbalance of women to men especially in the late adulthood and senescence phases, but this is undoubtably a subject worth investigating for differences between sexes in future lifespan work.

In conclusion these results describe the relationship between brain microstructure and age and illustrate how dMRI correlates of cellular microstructure changes throughout the lifespan. These results provide a powerful illustration of the dramatic shifts in microstructure metrics during development and aging and highlight the potential of 3T-CSD as a tool to study development and aging. Establishing patterns of microstructure change across the lifespan will aid in studying and interpreting microstructure change in the presence of pathology. Studies analyzing dMRI microstructure in developing or aging cohorts should address shifts in anisotropic and isotropic signal in development and dramatically increased levels of extracellular water in aging.

